# Comprehensive Glycoproteomic Analysis of Chinese Hamster Ovary Cells

**DOI:** 10.1101/318865

**Authors:** Ganglong Yang, Yingwei Hu, Shisheng Sun, Chuanzi Ouyang, Weiming Yang, Michael Betenbaugh, Hui Zhang

## Abstract

The Chinese hamster ovary (CHO) cell line is a major expression system for the production of therapeutic proteins, the majority of which are glycoproteins, such as antibodies and erythropoietin (EPO). The characterization of the glycosylation profiles is critical to understand the important role of glycosylation on therapeutic glycoproteins from CHO cells. In this study, a large scale glycoproteomic workflow was established and applied to CHO-K1 cells expressing EPO. The workflow includes enrichment of intact glycopeptides from CHO-K1 cell lysate and medium using hydrophilic enrichment, fractionation of the obtained intact glycopeptides (IGPs) by basic reversed phase liquid chromatography (bRPLC), analyzing the glycopeptides using LC-MS/MS, and annotating the results by GPQuest 2.0. A total of 10,338 N-linked glycosite-containing IGPs were identified, representing 1,162 unique glycosites in 530 glycoproteins, including 71 unique atypical N-linked IGPs on 18 atypical N-glycosylation sequons with an overrepresentation of the N-*X*-C motifs. Moreover, we compared the glycoproteins from CHO cell lysate with those from medium using the in-depth N-linked glycoproteome data. The obtained large scale glycoproteomic data from intact N-linked glycopeptides in this study is complementary to the genomic, proteomic, and N-linked glycomic data previously reported for CHO cells. Our method has the potential to accelerate the production of recombinant therapeutic glycoproteins.

## Introduction

The Chinese hamster ovary (CHO) cell line is the major expression system used for the efficient production of recombinant proteins, the majority of which are therapeutic glycoproteins, including erythropoietin (EPO), coagulation factors, and antibodies (Walsh 2014). A major focus of current glycosylation engineering efforts is to produce therapeutic glycoproteins with optimal yield and human-like post-translational modifications (PTMs). Glyco-engineered CHO cells have been comprehensively investigated by different biological, analytical, and molecular engineering approaches, including genomics (Birzele et al. 2010; Jones et al. 2010; Xu et al. 2011; Lewis et al. 2013; Chung et al. 2015), proteomics (Baycin-Hizal et al. 2012), glycoproteomics (Yang et al. 2014), and glycomics (North et al. 2010; Yang et al. 2015c), to understand the final characteristics of the recombinant therapeutic glycoproteins. These studies aim to optimize therapeutic glycoprotein drug production, thereby improving therapeutic efficacy and ultimately reducing side effects, toxicity, and cost in the pharmaceutical industry.

As one of the most prevalent protein modifications, N-linked glycosylation plays a vital role in many biological process, especially in protein folding, cell adhesion, cell-matrix interactions, cellular signaling, intracellular/extracellular targeting to organelles and pathogenesis of different diseases (Cummings and Pierce 2014; Xu and Ng 2015). The complexity of protein glycosylation, including multiple glycosites within a glycoprotein (macro-heterogeneity), site occupancy and different glycan structures at each glycosite (micro-heterogeneity), poses great challenges for a comprehensive analysis of glycoproteins expressed in a cell or organism to investigate the structural and functional role of glycans within glycoproteins. A number of methods have been used to characterize glycoproteins, including (i) glycosite identification using approaches such as Solid-Phase Extraction of Glycosite-containing peptides (SPEG) (Zhang et al. 2003), lectin enrichment (Kaji et al. 2003; Zielinska et al. 2010) and hydrophilic enrichment (Wada et al. 2004); (ii) glycome profiling, such as total native glycans (Fujitani et al. 2013), sialic acid derivatization (Shah et al. 2013; de Haan et al. 2015; Kammeijer et al. 2017; Yang et al. 2017a), reducing end labeling through stable isotopic labeling (Bigge et al. 1995; Ruhaak et al. 2010; Yang et al. 2015a; Yang et al. 2015b), and permethylation (Ciucanu and Costello 2003; Kang et al. 2005; Shubhakar et al. 2016); (iii) site-specific intact glycopeptides (IGPs) analysis (Scott et al. 2011; Parker et al. 2013; Khatri et al. 2016) using methods such as N-linked Glycan And Glycosite (NGAG) (Sun et al. 2016) analysis, Tool for Rapid Analysis of glycopeptide by Permethylation (TRAP) (Shajahan et al. 2017), Electron-Transfer/Higher-Energy Collision Dissociation (EThcD) mass spectrometry (Yu et al. 2017), and Hydrophilic Interaction Liquid Chromatography (HILIC) enrichment (Takegawa et al. 2006; Zhang et al. 2013). Site-specific IGP analysis is widely considered to be the most promising strategy to comprehensively characterize glycoproteins, but the IGP analysis workflow including enrichment method, mass spectrometric analysis and annotated software still needs to be developed to increase the glycoproteome coverage and precision. Recently, highly-comprehensive methods for IGP enrichment using HILIC-based methods were compared in our lab, and mixed anion exchange (MAX) extraction cartridges were selected as the optimal enrichment method for intact N-linked glycopeptides (Yang et al. 2017b). In addition, 2D fractionation methods, including bRPLC, gel electrophoresis (SDS-PAGE), or strong cation exchange (SCX-HPLC) are very helpful to further improve the coverage of peptides or glycopeptides (Dowell et al. 2008; Wang et al. 2011; Jia et al. 2016).

Glycosylation of therapeutic glycoproteins cannot be directly predicted by the genomic data. The expression of glycoproteins, glycoprotein biosynthesis and constituents are dictated by the availability of nucleotide sugar synthesis, nucleotide sugar transporters, enzyme activities, and other cellular status, which lead to broad structural diversity. Structural and functional analyses showed that protein glycosylation, especially for bisecting N-acetylglucosamine, fucosylation and sialylation, substantially impacts the functional activities and circulatory half-life of therapeutic glycoproteins (Walsh and Jefferis 2006; Hossler et al. 2009). Precise and comprehensive analysis of site-specific IGPs is critical to understand and control the glycosylation of glycoproteins produced in CHO cell expression systems.

In this study, we established a workflow for the large-scale characterization of intact N-linked glycopeptides using (1) MAX enrichment, (2) bRPLC fractionation, (3) mass spectrometry analysis using Q-Exactive instrumentation, and (4) data analysis and annotation in GPQuest 2.0 (Toghi Eshghi et al. 2015; Hu et al. 2018). Using this method, we identified a total of 10,338 unique N-linked glycosite-containing IGPs, representing 1,162 unique glycosites from 530 glycoproteins in human erythropoietin (EPO)-expressing CHO-K1 cell lysate and medium. From this study, 71 unique N-linked atypical IGPs were also identified, representing 18 unique glycosites with overrepresentation of the atypical sequon, N-*X*-C.

## Results and Discussion

### Identification of N-linked glycosite-containing peptides

The majority of therapeutic glycoproteins secreted from CHO cells were found to be heavily modified by N-linked glycosylation (Xu et al. 2011). Assessment of protein glycosylation in CHO cells is very important for understanding the quality of CHO-derived glycoproteins. Using multiple separation and analytical methodologies is helpful to expand the number of enriched glycopeptides and identified glycoproteins. In this study, we first identified the N-linked glycosylation sites and glycoproteins by PNGase F digestion. The tryptic peptides were fractionated by basic RPLC, then glycopeptides in each fraction were enriched by hydrophilic MAX extraction. After de-glycosylation by PNGase F, the glycosite-containing peptides were analyzed by LC-MS/MS and identified by MS-GF+ (Granholm et al. 2014; Kim and Pevzner 2014) (Figure 1a). The assigned N-glycosite-containing peptides were filtered with 0.1% FDR at the glyco-Peptide Spectrum Match (PSM) level, 0.3% FDR at the peptide level, and 1.1% at the protein level with a 2 PSM requirement for each peptide. A total of 68,148 PSMs were identified from CHO-K1 cells, representing 4,549 unique N-linked glycosite-containing peptides from 2,276 proteins (Supplemental_Table S1).

**Figure 1.**
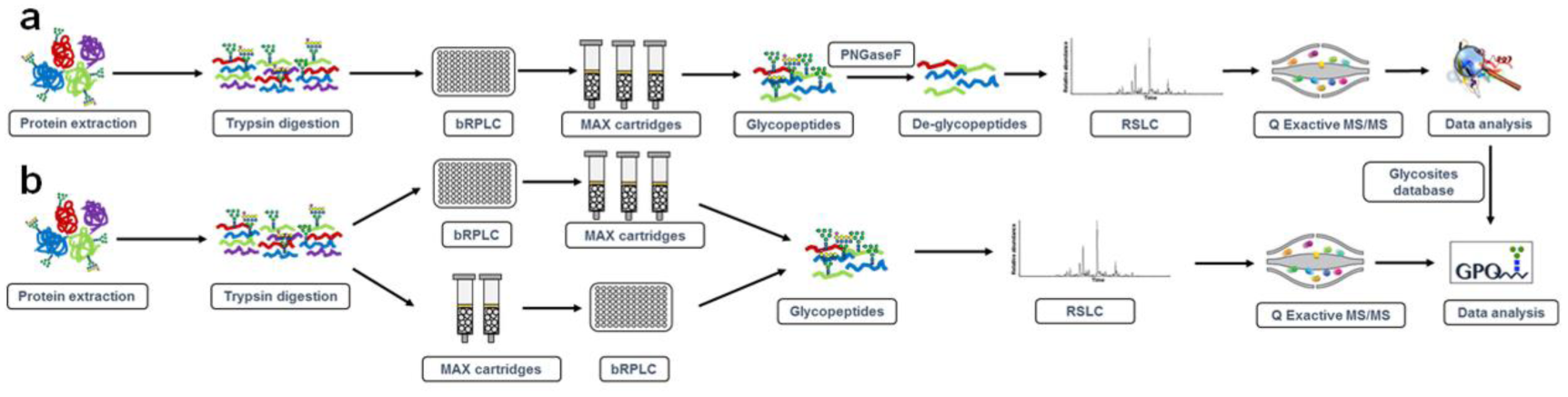
Workflows of intact glycopeptide analysis strategies for lysate and medium from human EPO-expressing CHO-K1 cells. (a) The workflow of large scale de-glycosylated peptide analysis enriched by MAX extraction cartridges. The proteins were digested, fractionated and MAX enriched. The glycopeptides were then de-glycosylated and analyzed by LC-MS/MS. (b) Different glycopeptide analysis strategies using MAX cartridge enrichment followed by fractionation or fractionation of global peptides followed by enrichment of intact glycopeptides using MAX cartridges for large-scale intact glycopeptides analysis.

### Intact N-linked glycopeptide analysis using different analytical workflows

To characterize the IGPs of the human EPO-expressing CHO-K1 cell line, different workflows were used in this study: (i) single LC-MS/MS analysis of enriched intact glycopeptides; (ii) direct analysis of global peptides using single LC-MS/MS or fractionation of global peptides by bRPLC followed by LC-MS/MS analysis; (iii) fractionation of global peptides using bRPLC followed by hydrophilic enrichment of intact glycopeptides from each fraction for LC-MS/MS analysis; and (iv) hydrophilic enrichment of intact glycopeptides followed by fractionation of enriched glycopeptides using bRPLC for LC-MS/MS analysis (Figure 1b). The prepared peptides were analyzed by a Q-Exactive mass spectrometer and annotated by GPQuest 2.0 using the glycosite-containing peptides described in the preceding section (Supplemental_Table S1) as the database. The intact N-linked glycopeptides were filtered using a 1% false discovery rate (FDR).

For single LC-MS/MS analysis of non-enriched and enriched intact glycopeptides, intact glycopeptides enriched from 1 μg or 100 μg of CHO cell global peptides by hydrophilic MAX columns were analyzed by LC-MS/MS and glycopeptides were identified by GPQuest 2.0. Five and 950 PSMs of intact glycopeptides were identified from 1 μg and 100 μg of CHO cell global peptides, respectively (Supplemental_Figure S1). More than a 190-fold increase of PSM identification was achieved when the initial peptide amount was increased from 1 μg to 100 μg for intact glycopeptide enrichment. We then compared the direct analysis of global peptides with or without fractionation using bRPLC followed by LC-MS/MS analysis. One microgram of global peptides were analyzed by single LC-MS/MS analysis or 100 μg global peptides were fractionated to 25 fractions and each fraction was analyzed by LC-MS/MS analysis. A total of 120 (from 1 μg) and 1,442 (from 100 μg) PSMs of intact glycopeptides were identified, respectively (Supplemental_Figure S1). The inclusion of the bRPLC separation resulted in a greatly (> 12-fold) increased number of identified PSMs. For large-scale identification of intact glycopeptides, (i) fractionation of peptides using bRPLC followed by MAX enrichment of intact glycopeptides from each fraction and (ii) MAX enrichment of intact glycopeptides followed by fractionation of enriched glycopeptides using bRPLC were established. Using a method entailing MAX enrichment followed by bRPLC (25 fractions), we were able to identify a total of 43,193 PSMs from 3.5 mg of global peptides, while 27,998 PSMs from an equal starting peptide amount were identified when the workflow started from the bRPLC fractionation step followed by MAX enrichment. There was a > 1.5-fold increase in PSM in the analysis of MAX-enriched intact glycopeptides followed by bRPLC strategy compared to the reverse method (Supplemental_Figure S1).

### Analysis of the N-linked IGPs in the human erythropoietin (EPO)-expressing CHO-K1 cell lysate and medium

Using CHO-K1 cell lysate, 8,391 unique N-linked glycosite-containing IGPs were identified, which matched with 1,090 glycosites, 507 glycoproteins, and 187 glycan compositions. In addition, the established large-scale IGPs analysis workflow was applied to the characterization of intact glycopeptides from secreted proteins in CHO-K1 cell culture. A total of 7,966 unique N-linked IGPs were identified in the medium of the CHO-K1 cells, which matched with 1,041 glycosites, 485 glycoproteins, and 171 glycan structures. In total, 10,338 unique N-linked glycosite-containing IGPs were identified from EPO-expressing CHO-K1 cell lysate and culture medium, matching to 1,162 glycosites, 530 glycoproteins, and 209 glycan compositions (Supplemental_Table S2 and Supplemental_Figure S2).

It is well known that most glycoproteins are extracellular proteins (transmembrane or secreted proteins) (Tian et al. 2010). Therefore, we predicted the subcellular location of identified glycoproteins using SignalP 4.1 (Nielsen 2017) and TMHMM 2.0 (Krogh et al. 2001) programs. Cell surface, secreted and transmembrane proteins were deemed as glycoproteins. The annotated signal peptides, protein glycosylation and Gene Ontology (GO) were evaluated in Uniprot. From the results, we found that 511 out of 530 (96.4%) identified proteins were highly likely to be glycoproteins, each of which contained either signal peptides, transmembrane helices, or reported glycosites (Supplemental_Table S3). These findings indicated the high specificity of our intact glycopeptide analysis workflow and the data analysis strategy towards comprehensive glycoproteomics characterization. The GO analysis also indicated that most of the glycoproteins were located at the cell surface or secreted to the extracellular space (Supplemental_Table S3). In addition, the molecular function of the identified glycoproteins was either binding or catalytic activity (Supplemental_Table S3).

To investigate the N-linked glycans from CHO cell lysate and medium, we characterized the IGPs by N-linked glycan classification. The identified 10,338 IGPs from both cell lysate and medium consisted of 4,452 high-mannose type glycan-containing glycopeptides and 5,886 complex and/or hybrid type glycan-containing glycopeptides. In these complex/hybrid glycan-containing glycopeptides, 3,941 and 1,471 were fucosylated and sialylated, respectively. High-mannose IGPs were the dominant type of glycopeptides (43.1%) and fucosylated IGPs were also very prevalent (38.1%), while sialylated glycopeptides were under represented (14.2%) among complex/hybrid IGPs in CHO cells (Figure 2a). The distribution of identified intact glycopeptides, glycoproteins and glycosites indicated that approximately 58.2% of the intact glycopeptides, 87.2% of the glycoproteins, and 83.5% of the glycosites were identified in both the cell lysate and the medium (Figure 2a and 2b). These findings suggest that most of the glycoproteins are common in cells and medium. However, the detailed structure of N-linked glycans on the glycoprotein differs between CHO cell lysate and medium. As an important component of glycoproteins, N-glycans and their site-specificity have not been thoroughly investigated in previously reported studies (Hart and Copeland 2010). In this study, glycans were assigned at their specific glycosylation sites through IGPs identification. Overall, 209 glycan structures were identified from human EPO-expressing CHO-K1 cell lysate and medium. Seven N-linked high-mannose glycans were identified in both cell and medium. For complex or hybrid glycans, different glycans were identified in the cell and the medium. The results showed that 78.7% complex/hybrid N-glycans were modified with fucose. Approximately 38.4% of the fucosylated N-linked glycans contained a core Fuc fragment ion (peptide+HexNAcFuc) that was present in the MS/MS spectra, indicating the core-fucosylated glycans (Figure 2c). In these N-linked fucosylated glycans, mono-fucosylated structures (46.8%) were the main type, while bi-fucosylated (13.7%), tri-fucosylated (15.1%) and multi-fucosylated (24.5%) were also widely identified (Figure 2c). In comparison, the majority of the N-linked sialylated glycans were mono-sialylated (54.2%) or bi-sialylated (28.9%) glycans. Only 16.9% of them were identified as multi-sialylated glycans (Figure 2c).

**Figure 2.**
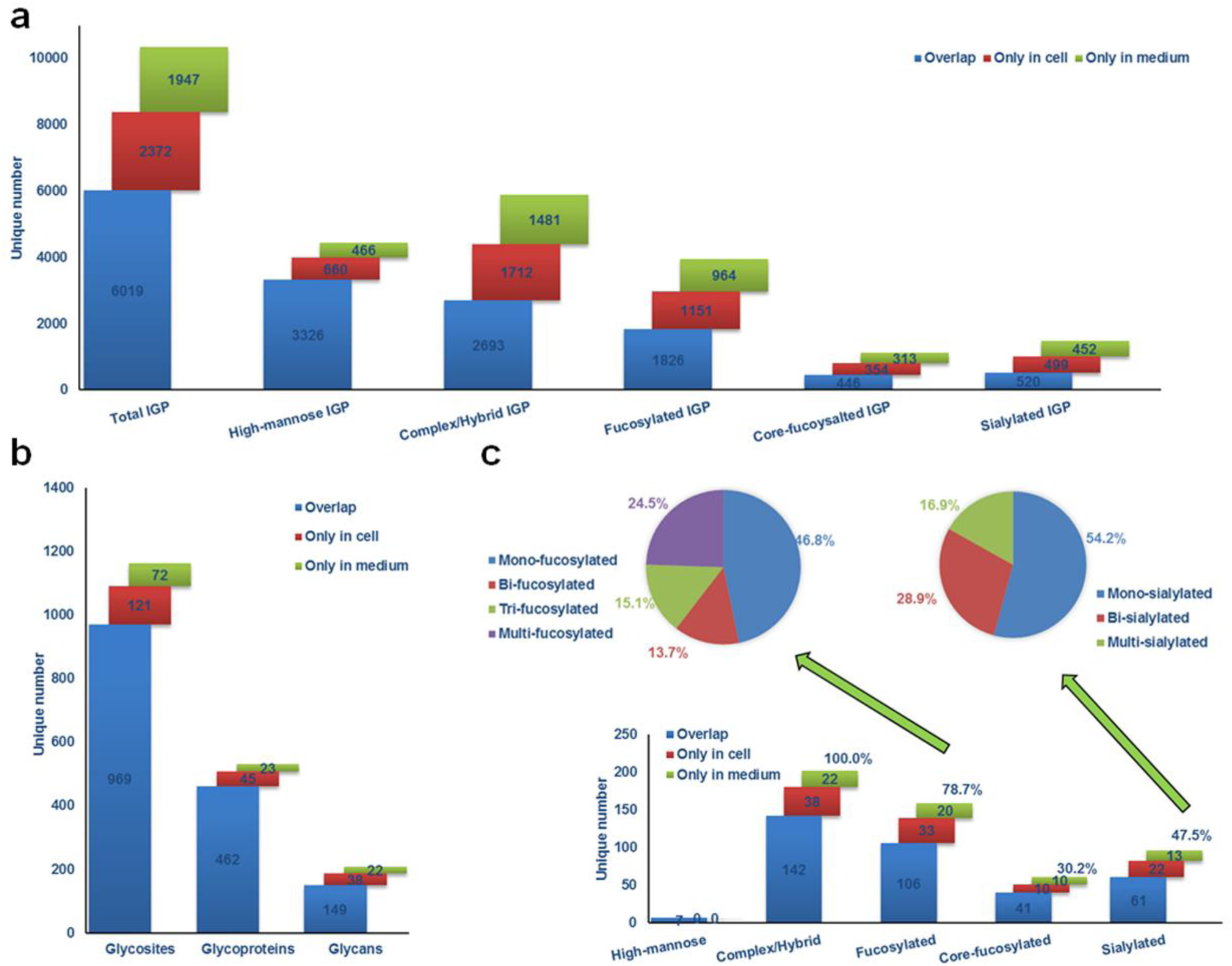
Depth of the identified intact glycopeptides in cell lysate and medium from human EPO-expressing CHO-K1 cells. (a) Identification and distribution of total and sub-type intact glycopeptides identified from cells and medium. (b) Distribution of identified glycosites and glycoproteins in cell lysate and medium, showing that most glycoproteins are present in CHO-K1 cell lysate and medium. (c) Identification and distribution of glycans and the composition and distribution of fucosylated and sialylated N-linked glycans in cell lysate and medium.

### Glycoprotein heterogeneity in CHO cells

Glycosylation heterogeneity is a common feature of glycoproteins. Each protein can potentially be glycosylated at multiple glycosylation sites and each glycosylation site can be modified by different glycans. Using our large-scale N-linked glycoproteomic data, we are able to reveal the macro-heterogeneity and micro-heterogeneity of glycoproteins in CHO cell lysate and medium. In the 530 N-linked glycoproteins identified from the intact glycoproteomic analysis, approximately 51.9% of the glycoproteins were identified with one N-linked glycosite, 24.2% of them were detected with two N-linked glycosites, and 10.4% with three N-linked glycosites. We identified a total of 55 (8.7%) glycoproteins that contained at least five glycosites (Figure 3a). The average number of N-linked glycosites was 2.2 per glycoprotein. To evaluate the relationship between the abundance of the glycoproteins and the index of protein glycosylation, we analyzed average protein abundance with the glycosite number on the same protein. We found that there is a positive relationship between the glycoprotein abundance and the index of its glycosylation with a P-value of 1.28E^-3^, indicating that the number of glycosites per protein is directly correlated with protein abundance, and additional glycosylation sites could be present in the low abundance glycoproteins and identified if an increased amount of protein was used for glycopeptide analysis (Supplemental_Figure S3a). The 5 most heavily glycosylated glycoproteins were prolow-density lipoprotein receptor-related protein 1 (30 unique N-linked glycosites), laminin subunit alpha-5 (19 unique N-linked glycosites), cation-independent mannose-6-phosphate receptor (18 unique N-linked glycosites), nicastrin (12 unique N-linked glycosites) and integrin alpha-3 (8 unique N-linked glycosites).

**Figure 3.**
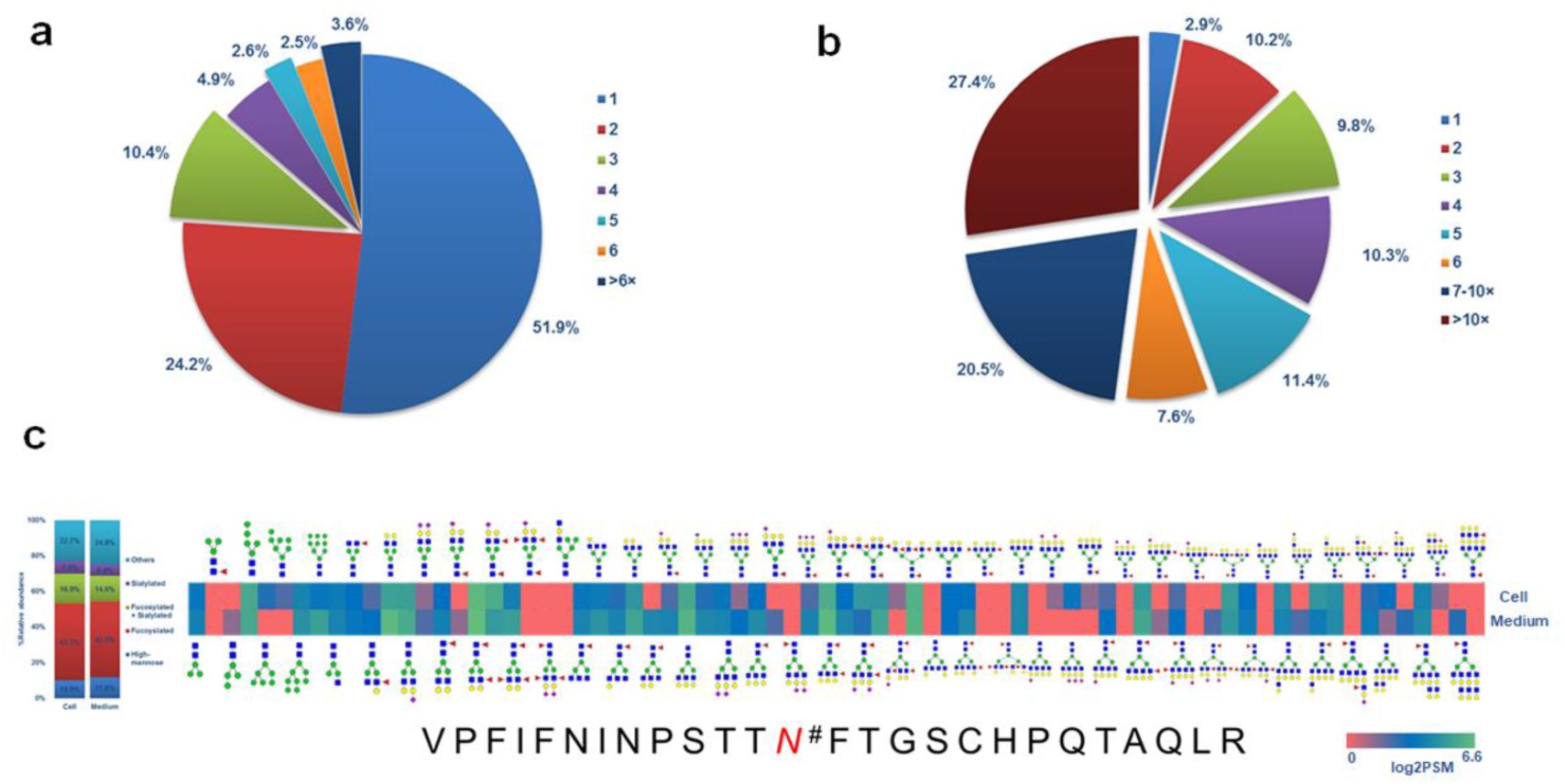
Heterogeneity of detected glycoproteins in CHO cell lysate and medium. (a) Distribution of glycosites per protein. (b) Distribution of glycans per glycosite. (c) Heat map of the differences in abundance of the sub-types of N-linked glycans between CHO cell lysate and medium on glycopeptide VPFIFNINPSTTN#FTGSCHPQTAQLR. # indicates an N-linked glycosite.

From the total of 1,162 N-linked glycosites, only 2.9% peptides carried a single glycan structure; approximately 10.2% and 9.8% glycosite-containing peptides were detected with two and three N-linked glycan structures at one glycosite, respectively. Notably, a group of 769 (66.9%) glycopeptides carried at least 5 N-linked glycan structures and 315 (27.4%) had at least 10 N-linked glycan structures at one glycosite (Figure 3b). The average number of N-linked glycan structures at one glycosite was 9.0. We also analyzed the variance of glycan structures on differentially abundant glycopeptides, finding a P-value of 2.06E^-20^ (Supplemental_Figure S3b), suggesting that the number of glycans identified from each glycosite is directly related to the abundance of the glycosylated peptides, indicating additional glycans could be present in the low abundance glycosites and could be identified if an increased amount of protein was used for glycopeptide analysis. In glycopeptide VPFIFNINPSTTN#FTGSCHPQTAQLR from Lysosome-associated membrane glycoprotein, we identified 74 N-linked glycan compositions. The most common glycan modification at this glycosite was found to be fucosylation (Figure 3c).

### Consensus motif preferences of N-linked glycosylation at canonical and atypical N-linked glycosites

The canonical N-linked glycopeptide sequon is known to be N-*X*-T/S (which *X* can be any amino acid except proline). In addition, some atypical N-glycosylation sequons have been discovered in recent years, such as N-*X*-C, N-*X*-V, and N-G-*X* (Zielinska et al. 2010; Sun and Zhang 2015). In the GPQuest 2.0 search engine, IGP identifications are determined based on the requirement of MS/MS spectra to contain oxonium ions and match to peptide or peptide+glycan fragment ions. Among the 10,338 identified MS/MS spectra, a total of 705 PSMs and 71 unique atypical N-linked IGPs were identified, which matched to 18 unique glycosites from 14 glycoproteins. Based on the spectra of atypical N-linked IGPs, two glycopeptides, TY<u>N#IV</u>LGSGQVVLGMDMGLR+N2H6F0S0G0 from glycoprotein peptidyl-prolyl cis-trans isomerase FKBP9 and LC<u>N#EC</u>SEGSFHLSK+N2H6F0S0G0 from basement membrane-specific heparin sulfate proteoglycan core protein (Figure 4a), were identified. The observed oxonium ions, peptides and peptide+glycan-related fragment ions from both peptides matched well.

**Figure 4.**
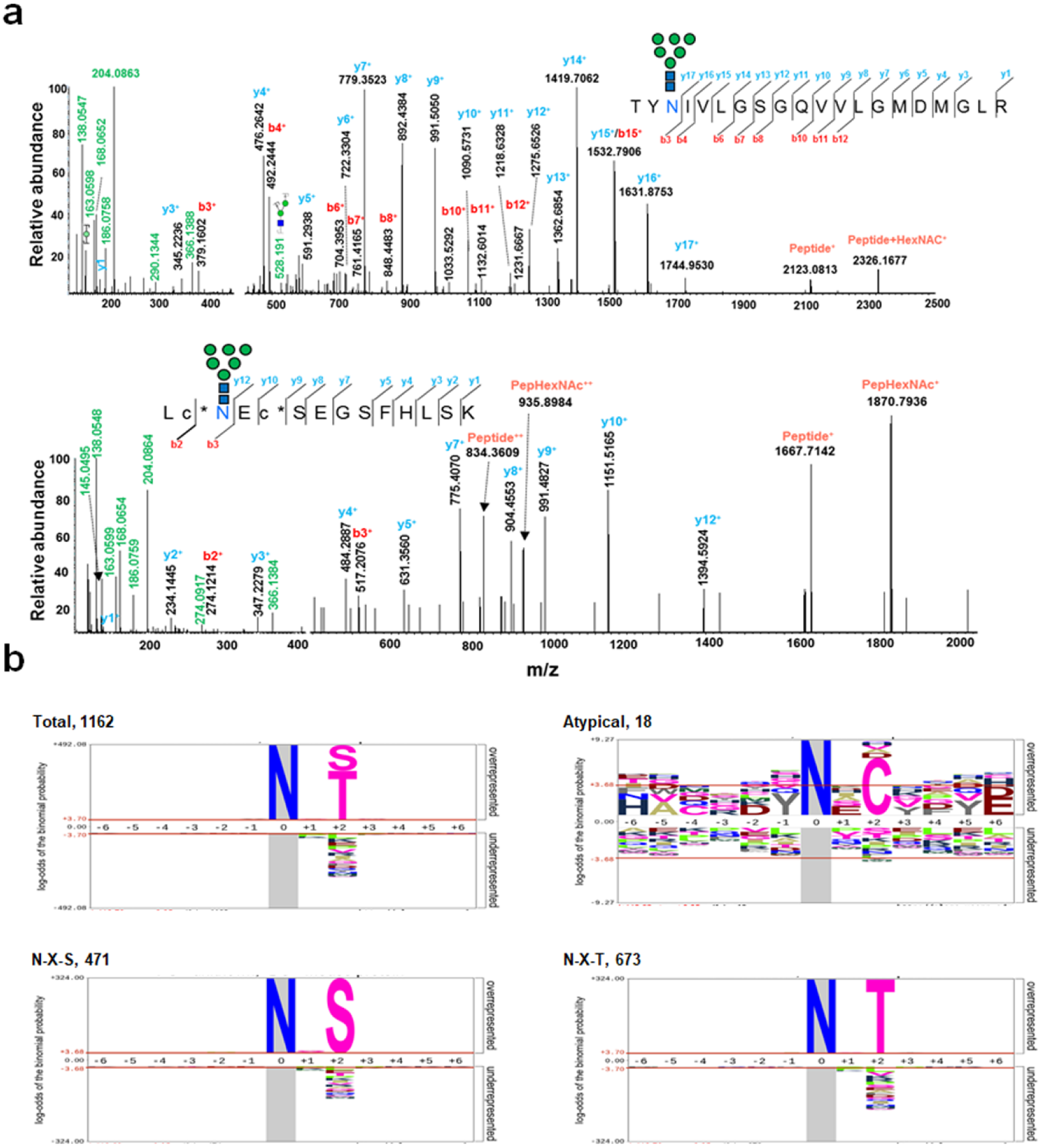
Identification of atypical N-linked glycopeptides and preference of N-glycosylation peptide consensus sequence. (a) Representative MS/MS spectra of the atypical N-glycopeptides of TYN#IVLGSGQVVLGMDMGLR+N2H6F0S0G0 from peptidyl-prolyl cis-trans isomerase FKBP9 and LCN#ECSEGSFHLSK+N2H6F0S0G0 from basement membrane-specific heparin sulfate proteoglycan core protein. # indicates an N-linked glycosite. (b) Distribution and preference of typical and atypical glycosite consensus sequence derived using pLogo.

To determine the preference of amino acids surrounding the canonical N-linked glycosylation sequons and discover other novel consensus sequons with our comprehensive data from this study, we compared the position-specific amino acid frequencies of the sequences (six amino acids from N-linked glycosylation sites at both termini) surrounding aspartic acid in both the canonical and the atypical N-linked glycopeptides using pLogo (O’Shea et al. 2013). A total of 1,162 unique glycosites were identified, of which 673 contained an N-X-T motif, 471 contained an N-X-S motif and 18 atypical glycosite motifs, and the consensus sequons of the identified glycosites were analyzed. As we examined the canonical sequons in the dataset, threonine was found to be more commonly present within the motif than serine at the +2 position (57.9% vs 40.5%). As expected, we found that proline was significantly underrepresented in all the scenarios at position +1 and +3. Of the three atypical N-glycosylation sequons identified (N-*X*-C, N-G-*X*, and N-*X*-V), only N-*X*-C was significantly overrepresented at 38.9% with a log-odds of binomial probabilities of +7.1 (Figure 4b).

### Relative abundance of N-linked IGPs between human EPO-expressing CHO-K1 cell lysate and medium

To investigate the abundance of N-linked IGPs differentially expressed between the CHO cell lysate and the medium, we performed a comparative analysis of IGPs using label-free quantification methods based on spectral counting. Overall, 43,742 and 62,665 PSMs were identified from the cell lysate and medium, respectively. A total of 6,763 and 7,966 unique IGPs were identified from cell lysate and medium, respectively. The relative abundance profiling and linear correlation analysis indicated that the N-glycosylation of proteins in cell lysate and medium was strongly correlated, as indicated by a Pearson correlation of 0.931, indicating that the overall glycosylation profiles of glycans in the glycosites between cell lysate and medium showed a statistical association (Figure 5a). However, we also found that the relative abundance of some IGPs were significantly altered between the cell lysate and the medium, such as EASQ<u>N^#^IT</u>YVCR-N2H6F0S0G0 with a log_2_PSM ratio of cell lysate/medium of 3.8, LQQEFHCCGS<u>N^#^NS</u>QDWR-N4H5F0S1G0 with a log_2_PSM ratio of cell/medium of 3.6; <u>N^#^ET</u>HSFCTACDESCK-N2H5F0S0G0 with a log_2_PSM ratio of cell/medium of 0.2, and LSPIHIAL<u>N^#^FS</u>LDPK-N2H6F0S0G0 with a log_2_PSM ratio of cell/medium of 0.2. Interestingly, these variabilities in N-glycosylation indicated that, while there was no significant difference in the global protein glycosylation, the stoichiometry could change dramatically at some of the N-glycosites, suggesting unique functions of these glycosylation events at different cellular locations.

**Figure 5.**
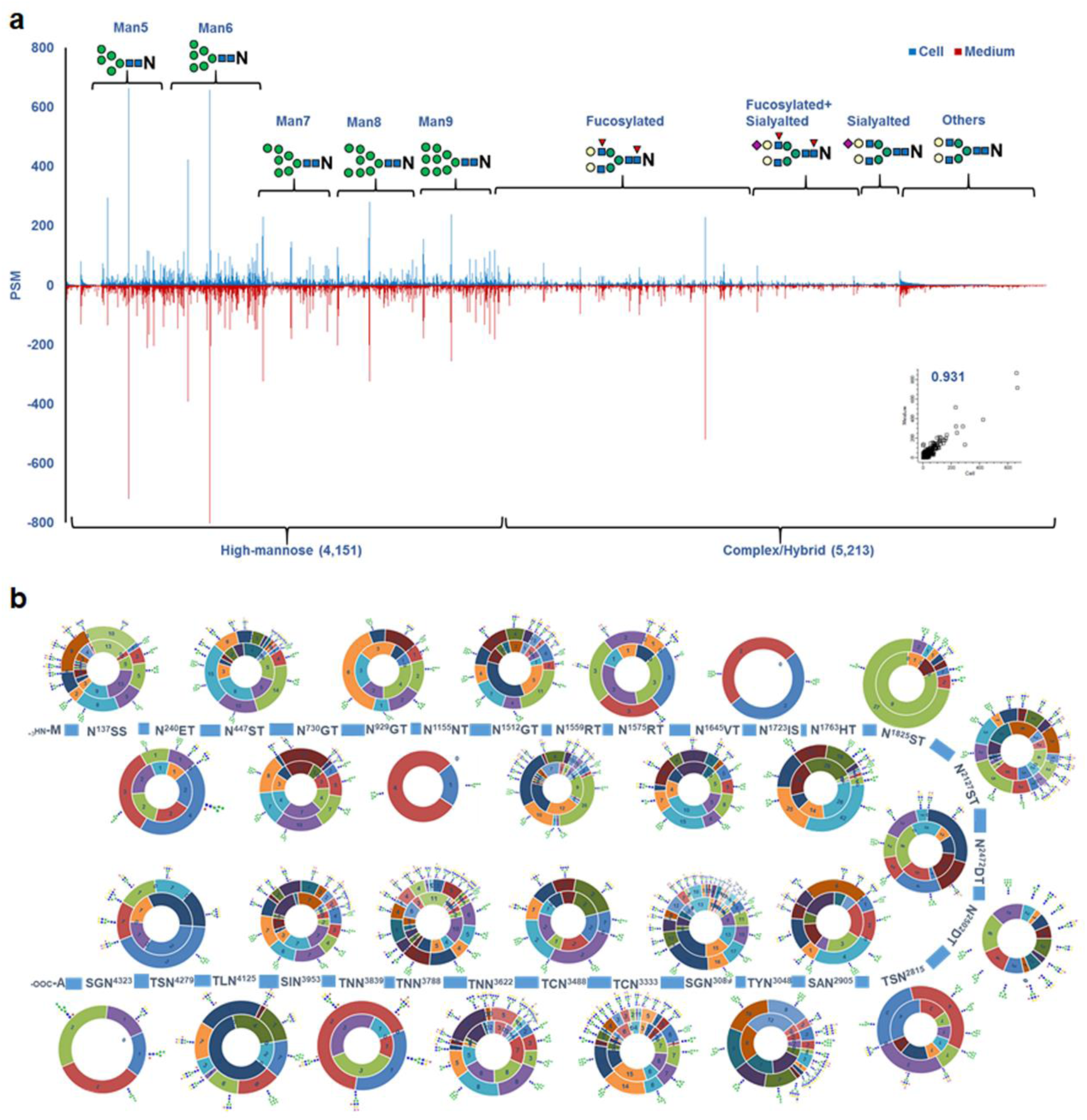
Relative abundance analysis of N-linked intact glycopeptides (IGPs). (a) Relative abundance profiling of IGPs between CHO cell lysate and medium, showing a strong correlation between the IGP abundance in cell lysate and medium. (b) The N-glycosylation of prolow-density lipoprotein receptor-related protein 1 in cell lysate and medium, including the structure and relative abundance (PSM) of glycans on each glycosite. The inner circle shows the relative abundance in CHO cell lysate and the outer circle shows the relative abundance in medium.

Most importantly, through the use of in-depth glycoproteomic profiling we are able to present a map of the protein N-glycosylation patterns in CHO cell lysate and medium. As mentioned, prolow-density lipoprotein receptor-related protein 1 was the most heavily glycosylated protein with 29 N-glycosites. The N-glycan map of prolow-density lipoprotein receptor-related protein 1 with different glycosylation sites based on our findings is shown in Figure 5b indicating the high degree of N-glycan micro-heterogeneity. This map not only presents the structure of the glycans but also the relative abundance of glycans on the glycosites. It is readily observed from the map that N-linked high-mannose glycans were identified on nearly every glycosite, especially for Man5 and Man6. Some of the N-linked glycosylation sites were present only either the CHO-K1 cell lysate or medium. Glycosylation at N^1155^NT and N^2502^DT were identified only in the cell lysate, while N^1723^IS and N^4279^ST were only identified in the cell medium. We also noticed two trends for the glycosylation of neighbor glycosites: (i) the glycosylation is similar between neighboring glycosites compared to distal glycosites. For example, N^1512^GT and N^1559^RT were both abundant with high mannose and biantennary complex glycans. This could occur because the glycotransferases or glycosidases recognize and are able to access part of the protein regions and modify these glycosites. (ii) The micro-heterogeneity was decreased when the glycosites were close to each other (i.e. glycosites N^1723^IS, N^1763^HT and N^1825^IS) (Supplemental_Figure S4). This phenomenon could occur because the glycan structure and site hindrance render the glycotransferase or glycosidase less accessible to the neighboring glycosites. Our results reveal that high-mannose glycan structures were typical in low degree micro-heterogeneity glycosites, while fucosylated glycan structures were common in high degree micro-heterogeneity glycosites.

### N-linked glycosylation of EPO

EPO is a model pharmaceutical protein in the development of CHO-based bioprocesses and the metabolic engineering of CHO cells for improved protein production (Ley et al. 2015). EPO glycosylation is important for its pharmacological properties. There have been many efforts to characterize the glycosylation of EPO in CHO cells, but most of the work focused on glycomics profiling or glycosite identification (Stübiger et al. 2005). With our large-scale IGPs identification method, the N-linked glycans of EPO at specific glycosylation sites were extensively characterized. Owing to the limitation that the first and second glycosites cannot be cleavage by trypsin, only two glycosite-containing peptides from EPO were identified with one peptide containing 2 glycosites (EAE<u>N#IT</u>TGCAEHCSLNE<u>N#IT</u>VPDTK and GQALLV<u>N#S</u>SQPWEPLQLHVDK (Table 1). For the glycosite-containing peptide with two glycosites, glycans conjugated to each glycosite could not be assigned due to the limied number of fragment ions presented for each glycosite. For the third glycosite, we found that 62 glycans were assigned, and complex or hybrid glycans were more common than other forms in the third glycosite from EPO. Almost all the glycans presented on EPO were also detected on other glycoproteins in the CHO cells. Glycan structure, relative abundance, and distribution of the identified glycopeptide, GQALLV<u>N#S</u>SQPWEPLQLHVDK, are summarized in Supplemental_Figure S5, of which 12 glycans (inside the red frame) have been previously identified in EPO glycomics studies (Jensen et al. 2012; Yin et al. 2015).

**Table 1.** The N-linked glycosylation of Erythropoietin (EPO) in CHO cell lysate and medium

|  | EAEN<sup>#</sup>ITTGCAEHCSLNEN<sup>#</sup>ITVPDTK |  |  | GQALLVN<sup>#</sup>SSQPWEPLQLHVDK |  |  |
| --- | --- | --- | --- | --- | --- | --- |
|  | CHO cell lysate<br>(Unique<br>IGPs/PSMs) | CHO medium<br>(Unique<br>IGPs/PSMs) | Overlap<br>(Unique<br>IGPs/PSMs) | CHO cell lysate<br>(Unique<br>IGPs/PSMs) | CHO medium<br>(Unique<br>IGPs/PSMs) | Overlap<br>(Unique<br>IGPs/PSMs) |
| High-mannose intact glycopeptides | 5/25 | 4/11 | 4/NA | 8/116 | 9/183 | 8/NA |
| Complex/Hybrid intact glycopeptides | 11/35 | 4/13 | 4/NA | 41/149 | 47/226 | 33/NA |
| Fucosylated intact glycopeptides | 9/16 | 2/3 | 2/NA | 26/99 | 34/142 | 23/NA |
| Sialylated intact glycopeptides | 3/4 | 0/0 | 0/NA | 14/40 | 21/63 | 12/NA |
| Intact glycopeptides | 16/60 | 8/24 | 8/NA | 49/265 | 54/409 | 41/NA |

### Conclusions

In this study, we presented a large-scale analysis of intact N-linked glycopeptides identified from CHO cells using established glycoproteomic workflows involving the enrichment of intact glycopeptides by MAX extraction cartridges followed bRPLC with high-resolution mass spectrometry. These results demonstrate the feasibility of the comprehensive exploration of the N-linked glycosylation of CHO cell lysate and medium. We also note that the coverage of CHO cell glycosylation characterization could be extensively improved in the near future by employing methodologies such as multiple proteases or top-down mass spectrometry.

## Methods

### Human EPO-expressing CHO-K1 cell culture

Human EPO-expressing CHO-K1 cells and medium were prepared by the National Institutes of Health (NIH). Briefly, the codon-optimized human EPO gene was overexpressed in CHO-K1 cells. After clone selection, a stable EPO-expressing CHO-K1 cell line was generated and adapted to suspension cells in CD-CHO media with 5mM Glutamine. Cells were seeded at 0.4 million/ml and incubated for 3 days with shaking. After shaking the culture, both supernatants and cell pellets were collected. Cell pellets were washed twice with phosphate buffered saline (PBS, pH 7.4) buffer. Cells were then lysed directly with 8 M urea/1 M NH4HCO3 solution and briefly sonicated until the solutions were clear. Protein concentrations were determined by BCA protein assay reagent (Thermo Scientific, Fair Lawn, NJ).

### Protein digestion

Proteins from Human EPO-expressing CHO-K1 cells and medium denatured in 8 M urea/1 M NH_4_HCO_3_ buffer were reduced by 10 mM TCEP at 37 °C for 1 h and alkylated by 15 mM iodoacetamide at room temperature for 30 min in the dark. The solutions were diluted eight-fold with ddH_2_O. Then sequencing grade trypsin (protein: enzyme, 40:1, w/w; Promega, Madison, WI) was added to the samples and incubated at 37 °C overnight. The solutions were acidified by acetic acid with pH<3. Samples were centrifuged at 13, 000g for 10 min and the supernatant was cleaned by C18 solid-phase extraction. Peptides were eluted from the C18 column in 60% ACN/0.1% TFA, and the peptide concentrations were measured by BCA reagent.

### Enrichment of N-linked glycopeptides using MAX Extraction Cartridges

For enrichment of N-linked glycopeptides, peptides after C18 desalting were adjusted to 95% ACN (v/v) 1% TFA (v/v) prior to loading onto Oasis MAX extraction cartridges (Waters). Cartridges were sequentially conditioned in ACN three times, 100 mM triethylammonium acetate three times, water five times and finally 95% ACN (v/v) 1% TFA (v/v) five times. Samples were loaded twice. The non-glycopeptides were washed by 95% ACN (v/v) 1% TFA (v/v) three times. Finally, glycopeptides were eluted in 50% ACN (v/v) 0.1% TFA (v/v), dried in a speed-vac and desalted by C18 solid-phase extraction.

### Peptide fractionation by basic reversed-phase liquid chromatography (bRPLC)

Basic reversed phase liquid chromatography was used with extensive fractionation to reduce sample complexity and thus reduce the likelihood of glycopeptides being co-isolated and co-fragmented(Bekker-Jensen et al. 2017). IGPs (~100 μg) extracted by MAX extraction cartridges and global peptides (~4 mg) from Human EPO-expressing CHO-K1 cells and medium were injected by a 1220 Series HPLC (Agilent Technologies, Inc., CA) into a Zorbax Extend-C18 analytical column containing 1.8 μm particles at a flow rate of 0.2 ml/min for IGPs or 3.5 μm particles at a flow rate of 1 ml/min for global peptides. The mobile-phase A consisted of 10 mM ammonium formate (pH 10) and B consisted of 10 mM ammonium formate and 90% ACN (pH 10). Sample separation was accomplished using the following linear gradient: 0–2% B, 10 min; 2–8% B, 5 min; 8–35% B, 85 min; 35–95% B, 5 min; 95–95% B, 15 min. Peptides were detected at 215 nm and 96 fractions were collected along with the LC separation in a time-based mode from 16 to 112 min. The 96 fractions of tryptic peptide samples were concatenated into 24 fractions by combining fractions 1, 25, 49, 73; 2, 26, 50, 74; etc. The samples were then dried in a speed-vac and stored at −80 °C until LC-MS/MS analysis.

### Glycopeptide enrichment and deglycosylation

For analysis of deglycosylated peptides, the tryptic digested peptides were fractionated by offline bRPLC. The glycopeptides of 24 fractions were enriched by MAX Extraction Cartridges. The captured glycopeptides were then deglycosylated by PNGase F (New England BioLabs). The de-glycosylated peptides were de-salted using Nuctip C18 tips (Waters). The peptides were dried and stored at −80 °C until LC-MS/MS analysis.

### NanoLC−MS/MS Analysis

The IGPs and de-glycosylated peptides were subjected two LC-MS/MS run per sample (or fraction) on a Q-Exactive mass spectrometer (Thermo Fisher Scientific, Bremen, Germany). The sample were re-suspended with 3% ACN and 0.1% FA. The samples were first separated on a Dionex Ultimate 3000 RSLC nano system (Thermo Scientific) with a PepMap RSLC C18 column (75 μm × 50 cm, 2μm, Thermo Scientific) protected by an Acclaim PepMap C18 column (100 μm × 2 cm, 5×m, Thermo Scientific). For the IGPs, the mobile phase consisted of 0.1% FA and 3% ACN in water (A) and 0.1% FA, 90% ACN (B) using a gradient elution of 0-2% B, 1 min; 2–8% B, 9 min; 8–31% B, 80 min, 31–38% B, 20 min; 38–95% B, 5 min; 95% B, 10 min; 95–2% B, 4 min. The flow rate was kept at 0.300 μL/min. Data-dependent higher-energy collisional dissociation (HCD) fragmentation was performed on the 12 most abundant ions. The spray voltage was set to 1.5 kV. Spectra (AGC target 3 × 10^6^ and maximum injection time 60 ms) were collected from 400–2000 m/z at a resolution of 70, 000 followed by data-dependent HCD MS/MS (at a resolution of 35, 000, NCE 32%, intensity threshold of 4.2 × 10^4^, AGC target 2 × 10^5^ and maximum injection time 120 ms) of the ions using an isolation window of 1.4 m/z. Charge state screening was enabled to reject singly charged ions and ions with more than eight charges. A dynamic exclusion time of 30 sec was used to discriminate against previously selected ions. The de-glycosylated peptides were separated with gradient elution of 0-2% B, 1 min; 2–7% B, 9 min; 7–27% B, 80 min, 27–35% B, 22 min; 35–95% B, 3 min; 95% B, 10 min; 95–2% B, 6 min. Orbitrap MS1 spectra (AGC 3 × 10^6^) were collected from 400–1,800 m/z at a resolution of 70, 000 followed by data-dependent HCD MS/MS (resolution 3, 500, NCE 29%, intensity threshold of 3.3 × 10^4^, AGC target 2 × 10^5^ and maximum injection time 120 ms) of the 12 most abundant ions using an isolation width of 1.4 Da. Charge state screening was enabled to reject singly charged ions and ions with more than eight charges. A dynamic exclusion time of 30 s was used to discriminate against previously selected ions. For tryptic peptides, 110 m/z was set as the fixed first mass in MS/MS fragmentation to include all oxonium ions of glycopeptides.

### Data Analysis

For de-glycosylated peptides, acquired MS/MS spectra were searched using MS-GF+ against the RefSeq Cricetulus griseus protein database downloaded from the NCBI website with last update June 01, 2016, which contained 46,402 proteins. Database search parameters were set as follows: a maximum of two missed cleavage sites permitted for trypsin digestion, 10 ppm precursor mass tolerance, 0.06 Da fragment mass tolerance, carbamidomethylation (C, +57.0215 Da) as a fixed modification, and oxidization (M, +15.9949 Da) and deamidation (N, +0.98 Da) as dynamic modifications. The results were filtered with 1% FDR for PSM and 2 PSMs were required for a peptide. Spectral counting was used to quantify the peptides identified from LC-MS/MS data.

For IGPs identification, the data were searched using an in-house developed glycopeptide analysis software GPQuest 2.0, based on GPQuest(Toghi Eshghi et al. 2015). The database of glycosites were the glycosite-containing peptide data from de-glycopeptide methods. The human erythropoietin fasta protein sequence was also added to the protein database. The Cricetulus griseus glycan database was from the previous CHO cell glycomics profiling studys(North et al. 2010). The databases contains 57,653 predicted glycosites and 343 glycan structure entries. The parameters for mass tolerance of MS1 and MS2 were 10 ppm and 20 ppm, respectively. The spectra containing an oxonium ion *m/z* 204.09 were chosen for further searching. Results were filtered based on the following criteria: (1) false discovery rate (FDR) less than 1%, (2) ≥3 PSMs for each peptide were required, (3) all the PSMs should be annotated by at least one N-linked glycans, and (4) for the atypical glycoprotein, at least one typical N-linked glycosite-containing peptide was required.

### Data access

The raw data of mass spectrometry from this study will be submitted. All the annotated intact glycopeptides and deglycopeptides are listed in Supplemental tables S1-S3.

## Acknowledgements

This work was supported by the National Heart, Lung and Blood Institute, Programs of Excellence in Glycosciences (PEG, Grant P01HL107153), National Cancer Institute, the Clinical Proteomic Tumor Analysis Consortium (CPTAC, Grant U24CA210985) and the Early Detection Research Network (EDRN, U01CA152813), and National Science Foundation (Grant 124230). The authors declare no competing conflicts of interest.

## Author contributions

HZ and GY designed the experiments and wrote and reviewed the manuscript. GY, SS and CO performed the cell culture and preparation, intact glycopeptide enrichment and mass spectrometry analysis. YH and WY contributed to prepare the software and analyze the data.

## Supplemental Figures and Tables

Table S1: The glycosite-containing peptides identified from CHO cells.

Table S2: Identification of intact glycopeptides from CHO-K1 cell lysate and medium.

Table S3: Subcellular location, signal peptides, and transmembrane segments for identified glycoproteins from CHO cell lysate and medium.

Figure S1: Identification of Peptide Spectrum Matches (PSMs) for intact glycopeptides from human EPO-expressing CHO-K1 cell lysate using different analytical workflows.

Figure S2: Identification of N-linked intact glycopeptides from CHO-K1 cell lysate and medium using MAX enrichment of intact glycopeptides followed by fractionation of enriched glycopeptides using bRPLC.

Figure S3: Relationship between the number of N-linked glycosylation sites and protein abundance in CHO cell lysate and medium. (a) Distribution of average abundance of glycoproteins with different numbers of glycosites represented by violin plots. (b) Distribution of average abundance of glycopeptides with different numbers of glycans represented by boxplots.

Figure S4: Proportion of identified IGPs with different N-glycan compositions on prolow-density lipoprotein receptor-related protein 1 from CHO cell lysate and medium.

Figure S5. Relative abundance of specific glycan-containing forms of the glycosite-containing EPO peptide, GQALLV*<u>N^#^</u>*<u>SS</u>QPWEPLQLHVDK from CHO cell lysate and medium.

## References

Baycin-Hizal D, Tabb DL, Chaerkady R, Chen L, Lewis NE, Nagarajan H, Sarkaria V, Kumar A, Wolozny D, Colao J, et al. 2012. Proteomic analysis of Chinese hamster ovary cells. J Proteome Res 11: 5265–5276.

Bekker-Jensen DB, Kelstrup CD, Batth TS, Larsen SC, Haldrup C, Bramsen JB, Sorensen KD, Hoyer S, Orntoft TF, Andersen CL, et al. 2017. An optimized shotgun strategy for the rapid generation of comprehensive human proteomes. Cell Syst 4: 587–599.

Bigge JC, Patel TP, Bruce JA, Goulding PN, Charles SM, Parekh RB. 1995. Nonselective and efficient fluorescent labeling of glycans using 2-amino benzamide and anthranilic acid. Anal Biochem 230: 229–238.

Birzele F, Schaub J, Rust W, Clemens C, Baum P, Kaufmann H, Weith A, Schulz TW, Hildebrandt T. 2010. Into the unknown: expression profiling without genome sequence information in CHO by next generation sequencing. Nucleic Acids Res 38: 3999–4010.

Chung CY, Yin B, Wang Q, Chuang KY, Chu JH, Betenbaugh MJ. 2015. Assessment of the coordinated role of ST3GAL3, ST3GAL4 and ST3GAL6 on the alpha2,3 sialylation linkage of mammalian glycoproteins. Biochem Biophys Res Commun 463: 211–215.

Ciucanu I, Costello CE. 2003. Elimination of oxidative degradation during the per-O-Methylation of carbohydrates. J AM CHEM SOC 125: 16213–16219.

Cummings RD, Pierce JM. 2014. The challenge and promise of glycomics. Chem Biol 21: 1–15.

de Haan N, Reiding KR, Haberger M, Reusch D, Falck D, Wuhrer M. 2015. Linkage-specific sialic acid derivatization for MALDI-TOF-MS profiling of IgG glycopeptides. Anal Chem 87: 8284–8291.

Dowell JA, Frost DC, Zhang J, Li L. 2008. Comparison of two-dimensional fractionation techniques for shotgun proteomics. Anal Chem 80: 6715–6723.

Fujitani N, Furukawa J, Araki K, Fujioka T, Takegawa Y, Piao J, Nishioka T, Tamura T, Nikaido T, Ito M, et al. 2013. Total cellular glycomics allows characterizing cells and streamlining the discovery process for cellular biomarkers. Proc Natl Acad Sci U S A 110: 2105–2110.

Granholm V, Kim S, Navarro JC, Sjolund E, Smith RD, Kall L. 2014. Fast and accurate database searches with MS-GF+Percolator. J Proteome Res 13: 890–897.

Hart GW, Copeland RJ. 2010. Glycomics hits the big time. Cell 143: 672–676.

Hossler P, Khattak SF, Li ZJ. 2009. Optimal and consistent protein glycosylation in mammalian cell culture. Glycobiology 19: 936–949.

Hu Y, Shah P, Clark DJ, Ao M, Zhang H. 2018. Reanalysis of global proteomic and phosphoproteomic data identified a large number of glycopeptides. Anal Chem. Revision.

Jensen PH, Karlsson NG, Kolarich D, Packer NH. 2012. Structural analysis of N- and O-glycans released from glycoproteins. Nat Protoc 7: 1299–1310.

Jia X, Chen J, Sun S, Yang W, Yang S, Shah P, Hoti N, Veltri B, Zhang H. 2016. Detection of aggressive prostate cancer associated glycoproteins in urine using glycoproteomics and mass spectrometry. Proteomics 16: 2989–2996.

Jones MB, Tomiya N, Betenbaugh MJ, Krag SS. 2010. Analysis and metabolic engineering of lipid-linked oligosaccharides in glycosylation-deficient CHO cells. Biochem Biophys Res Commun 395: 36–41.

Kaji H, Saito H, Yamauchi Y, Shinkawa T, Taoka M, Hirabayashi J, Kasai K-i, Takahashi N, Isobe T. 2003. Lectin affinity capture, isotope-coded tagging and mass spectrometry to identify N-linked glycoproteins. Nat Biotechnol 21: 667–672.

Kammeijer GSM, Jansen BC, Kohler I, Heemskerk AAM, Mayboroda OA, Hensbergen PJ, Schappler J, Wuhrer M. 2017. Sialic acid linkage differentiation of glycopeptides using capillary electrophoresis – electrospray ionization – mass spectrometry. Sci Rep 7, 10.1038/s41598-017-03838-y.

Kang P, Mechref Y, Klouckova I, Novotny MV. 2005. Solid-phase permethylation of glycans for mass spectrometric analysis. Rapid Commun Mass Spectrom 19: 3421–3428.

Khatri K, Klein JA, White MR, Grant OC, Leymarie N, Woods RJ, Hartshorn KL, Zaia J. 2016. Integrated Omics and computational glycobiology reveal structural basis for influenza A virus glycan microheterogeneity and host interactions. Mol Cell Proteomics 15: 1895–1912.

Kim S, Pevzner PA. 2014. MS-GF+ makes progress towards a universal database search tool for proteomics. Nat Commun 5: 5277, DOI: 10.1038/ncomms6277.

Krogh A, BjörnLarsson, Heijne G, L.LSonnhammer E. 2001. Predicting transmembrane protein topology with a hidden markov model application to complete genomes. J Mol Biol 305: 567–580.

Lewis NE, Liu X, Li Y, Nagarajan H, Yerganian G, O’Brien E, Bordbar A, Roth AM, Rosenbloom J, Bian C, et al. 2013. Genomic landscapes of Chinese hamster ovary cell lines as revealed by the Cricetulus griseus draft genome. Nat Biotechnol 31: 759–765.

Ley D, Seresht AK, Engmark M, Magdenoska O, Nielsen KF, Kildegaard HF, Andersen MR. 2015. Multi-omic profiling of EPO-producing Chinese hamster ovary cell panel reveals metabolic adaptation to heterologous protein production. Biotechnol Bioeng 112: 2373–2387.

Nielsen H. 2017. Predicting Secretory Proteins with SignalP. In Protein Function Prediction: Methods and Protocols, doi:10.1007/978-1-4939-7015-5_6 (ed. D Kihara), pp. 59-73. Springer New York, New York, NY.

North SJ, Huang HH, Sundaram S, Jang-Lee J, Etienne AT, Trollope A, Chalabi S, Dell A, Stanley P, Haslam SM. 2010. Glycomics profiling of Chinese hamster ovary cell glycosylation mutants reveals N-glycans of a novel size and complexity. J Biol Chem 285: 5759–5775.

O’Shea JP, Chou MF, Quader SA, Ryan JK, Church GM, Schwartz D. 2013. pLogo: a probabilistic approach to visualizing sequence motifs. Nat Methods 10: 1211–1212.

Parker BL, Thaysen-Andersen M, Solis N, Scott NE, Larsen MR, Graham ME, Packer NH, Cordwell SJ. 2013. Site-specific glycan-peptide analysis for determination of N-glycoproteome heterogeneity. J Proteome Res 12: 5791–5800.

Ruhaak LR, Steenvoorden E, Koeleman CA, Deelder AM, Wuhrer M. 2010. 2-picoline-borane: a non-toxic reducing agent for oligosaccharide labeling by reductive amination. Proteomics 10: 2330–2336.

Scott NE, Parker BL, Connolly AM, Paulech J, Edwards AV, Crossett B, Falconer L, Kolarich D, Djordjevic SP, Hojrup P, et al. 2011. Simultaneous glycan-peptide characterization using hydrophilic interaction chromatography and parallel fragmentation by CID, higher energy collisional dissociation, and electron transfer dissociation MS applied to the N-linked glycoproteome of Campylobacter jejuni. Mol Cell Proteomics 10: 10.1074/mcp.M000031-MCP000201.

Shah P, Yang S, Sun S, Aiyetan P, Yarema KJ, Zhang H. 2013. Mass spectrometric analysis of sialylated glycans with use of solid-phase labeling of sialic acids. Anal Chem 85: 3606–3613.

Shajahan A, Supekar NT, Heiss C, Ishihara M, Azadi P. 2017. Tool for Rapid Analysis of glycopeptide by Permethylation (TRAP) via one-pot site mapping and glycan analysis. Anal Chem 89: 10734–10743.

Shubhakar A, Kozak RP, Reiding KR, Royle L, Spencer DI, Fernandes DL, Wuhrer M. 2016. Automated high-throughput permethylation for glycosylation analysis of biologics using MALDI-TOF-MS. Anal Chem 88: 8562–8569.

Stübiger G, Marchetti M, Nagano M, Grimm R, Gmeiner G, Reichel C, Allmaier G. 2005. Characterization of N- and O-glycopeptides of recombinant human erythropoietins as potential biomarkers for doping analysis by means of microscale sample purification combined with MALDI-TOF and quadrupole IT/RTOF mass spectrometry. J Sep Sci 28: 1764–1778.

Sun S, Shah P, Eshghi ST, Yang W, Trikannad N, Yang S, Chen L, Aiyetan P, Hoti N, Zhang Z et al. 2016. Comprehensive analysis of protein glycosylation by solid-phase extraction of N-linked glycans and glycosite-containing peptides. Nat Biotechnol 34: 84–88.

Sun S, Zhang H. 2015. Identification and validation of atypical N-glycosylation sites. Anal Chem 87: 11948–11951.

Takegawa Y, Deguchi K, Keira T, Ito H, Nakagawa H, Nishimura S. 2006. Separation of isomeric 2-aminopyridine derivatized N-glycans and N-glycopeptides of human serum immunoglobulin G by using a zwitterionic type of hydrophilic-interaction chromatography. J Chromatogr A 1113: 177–181.

Tian Y, Kelly-Spratt KS, Kemp CJ, Zhang H. 2010. Mapping tissue-specific expression of extracellular proteins using systematic glycoproteomic analysis of different mouse tissues. J Proteome Res 9: 5837–5847.

Toghi Eshghi S, Shah P, Yang W, Li X, Zhang H. 2015. GPQuest: A spectral library matching algorithm for site-specific assignment of tandem mass spectra to intact N-glycopeptides. Anal Chem 87: 5181–5188.

Wada Y, Tajiri M, Yoshida S. 2004. Hydrophilic affinity isolation and MALDI multiple-stage tandem mass spectrometry of glycopeptides for glycoproteomics. Anal Chem 76: 6560–6565.

Walsh G. 2014. Biopharmaceutical benchmarks. 2014. Nat Biotechnol 32: 992–1002.

Walsh G, Jefferis R. 2006. Post-translational modifications in the context of therapeutic proteins. Nat Biotechnol 24: 1241–1252.

Wang Y, Yang F, Gritsenko MA, Wang Y, Clauss T, Liu T, Shen Y, Monroe ME, Lopez-Ferrer D, Reno T et al. 2011. Reversed-phase chromatography with multiple fraction concatenation strategy for proteome profiling of human MCF10A cells. Proteomics 11: 2019–2026.

Xu C, Ng DT. 2015. Glycosylation-directed quality control of protein folding. Nat Rev Mol Cell Biol 16: 742–752.

Xu X, Nagarajan H, Lewis NE, Pan S, Cai Z, Liu X, Chen W, Xie M, Wang W, Hammond S, et al. 2011. The genomic sequence of the Chinese hamster ovary (CHO)-K1 cell line. Nat Biotechnol 29: 735–741.

Yang G, Tan Z, Lu W, Guo J, Yu H, Yu J, Sun C, Qi X, Li Z, Guan F. 2015a. Quantitative glycome analysis of N-glycan patterns in bladder cancer vs normal bladder cells using an integrated strategy. J Proteome Res 14: 639–653.

Yang S, Wang M, Chen L, Yin B, Song G, Turko IV, Phinney KW, Betenbaugh MJ, Zhang H, Li S. 2015b. QUANTITY: An isobaric tag for quantitative glycomics. Sci Rep 5: 17585, DOI: 10.1038/srep17585.

Yang S, Zhang L, Thomas S, Hu Y, Li S, Cipollo J, Zhang H. 2017a. Modification of sialic acids on solid phase: accurate characterization of protein sialylation. Anal Chem 89: 6330–6335.

Yang W, Shah P, Hu Y, Toghi Eshghi S, Sun S, Liu Y, Zhang H. 2017b. Comparison of enrichment methods for intact N- and O-Linked glycopeptides using strong anion exchange and hydrophilic interaction liquid chromatography. Anal Chem 89: 11193–11197.

Yang Z, Halim A, Narimatsu Y, Josh HJ, Steentoft C, Schjoldager KT-BG, Schulz MA, Sealover NR, Kayser KJ, Bennett EP, et al. 2014. The GalNAc-type O-Glycoproteome of CHO cells characterized bu the simplecell strategy. Mol Cell Proteomics 13: 12.

Yang Z, Wang S, Halim A, Schulz MA, Frodin M, Rahman SH, Vester-Christensen MB, Behrens C, Kristensen C, Vakhrushev SY, et al. 2015. Engineered CHO cells for production of diverse, homogeneous glycoproteins. Nat Biotechnol 33: 842–844.

Yin B, Gao Y, Chung CY, Yang S, Blake E, Stuczynski MC, Tang J, Kildegaard HF, Andersen MR, Zhang H, et al. 2015. Glycoengineering of Chinese hamster ovary cells for enhanced erythropoietin N-glycan branching and sialylation. Biotechnol Bioeng 112: 2343–2351.

Yu Q, Wang B, Chen Z, Urabe G, MatthewS. Glover, Shi X, Guo L-W, Kent C, Li L. 2017. Electron-transfer/higher-energy collision dissociation (EThcD)-enabled intact glycopeptide/glycoproteome characterization. J Am Soc Mass Spectrom 28: 1751–1764.

Zhang H, Li X-j, Martin DB, Aebersold R. 2003. Identification and quantification of N-linked glycoproteins using hydrazide chemistry, stable isotope labeling and mass spectrometry. Nat Biotechnol 21: 660–666.

Zhang Z, Wu Z, Wirth MJ. 2013. Polyacrylamide brush layer for hydrophilic interaction liquid chromatography of intact glycoproteins. J Chromatogr A 1301: 156–161.

Zielinska DF, Gnad F, Wisniewski JR, Mann M. 2010. Precision mapping of an in vivo N-glycoproteome reveals rigid topological and sequence constraints. Cell 141: 897–907.

